# Establishment of a simplified preparation method for single-nucleus RNA-sequencing and its application to long-term frozen tumor tissues

**DOI:** 10.1101/2020.10.23.351809

**Authors:** Kati J. Ernst, Konstantin Okonechnikov, Josephine Bageritz, Jan-Philipp Mallm, Andrea Wittmann, Kendra K. Maaß, Svenja Leible, Michael Boutros, Stefan M. Pfister, Marc Zuckermann, David T.W. Jones

**Affiliations:** Hopp Children’s Cancer Center Heidelberg (KiTZ), Heidelberg, Germany; Pediatric Glioma Research Group, German Cancer Research Center (DKFZ), Heidelberg, Germany; Division of Pediatric Neuro-oncology, German Cancer Research Center (DKFZ), Heidelberg, Germany; Division of Signaling and Functional Genomics, German Cancer Research Center (DKFZ), Heidelberg, Germany; Single-Cell Open Lab, German Cancer Research Center (DKFZ), Heidelberg, Germany; Department of Pediatric Hematology and Oncology, University Hospital, Heidelberg, Germany

## Abstract

Recent advances allowing the genomic analysis of individual cells from a bulk population have provided intriguing new insights into areas such as developmental processes and tumor heterogeneity. Most approaches to date, however, rely on the availability of fresh surgical specimens, thereby dramatically reducing the ability to profile particularly rare tissue types. Pediatric central nervous system tumors – the leading cause of childhood cancer deaths – represent one such example, where often only frozen rather than native material is available. Due to an increasing need for advanced techniques to understand the heterogeneity of these tumors, we optimized a method to isolate intact nuclei from long-term frozen pediatric glioma tissues. We performed a technical comparison between different single nucleus RNA-sequencing (snRNA-seq) systems using a patient-derived xenograft model as a test sample. Further, we applied the established nucleus isolation method to analyze frozen primary tissue from two pediatric central nervous system tumors – one pilocytic astrocytoma and one glioblastoma – allowing the identification of distinct tumor cell populations and infiltrating microglia. The results show that our fast, simple and low-cost nuclear isolation protocol provides intact nuclei, which can be used in both droplet-based 3’ transcriptome amplification (10X Genomics) and plate-based whole transcriptome amplification (Fluidigm C1) single-cell sequencing platforms, thereby dramatically increasing the potential for application of such methods to rare entities.

## Introduction

Single-cell RNA sequencing (scRNA-seq) is an increasingly popular method for investigating properties of heterogeneous tissues, which cannot easily be addressed with conventional bulk tissue RNA sequencing. Specifically, it allows the assessment of tumor heterogeneity and the identification of diverse tumor and non-tumor cell populations in a tumor mass. Whole-cell extractions for traditional scRNA-seq, however, require viable fresh tissue, whereas often only frozen tumor biopsy samples are readily available.

The need for modern techniques that enable a thorough investigation and understanding of tumor biology is underlined by the clinical challenge presented by many tumor entities, such as childhood brain tumors. Central nervous system tumors are currently the leading cause of cancer-related morbidity and mortality in children in spite of all therapeutic efforts^1–4^. While there have been several studies looking at single-cell analysis of pediatric brain tumors from viable whole cells^5–8^, application to frozen tissue (which would open up a wealth of possible samples) has been limited to date. We therefore sought to develop a robust method for profiling of single nuclei (snRNA-seq) extracted from long-term frozen samples of pediatric brain tumors. Previous reports have indicated the utility of frozen material for snRNA-seq^9–11^, but brain tissues are especially problematic as starting material due to their high neuron composition and thus increased sensitivity to enzymatic cell dissociation^12^. We therefore focused on using only mild enzymatic buffers and a mechanical cell dissociation. We compared a density gradient nucleus isolation protocol (modified from Spalding et al 2005^13^ and Ernst et al 2014^14^) to other previously reported nucleus isolation methods, such as Nuclei EZ Prep (Sigma-Aldrich, NUC101-1KT), Isolation of Nuclei for Single-Cell RNA Sequencing (10X Genomics)^15^ and OptiPrep™^16^. None of these, however, resulted in an adequate sample quality (defined by a high yield of intact nuclei with minimal cell debris) when using the frozen brain tumor samples as starting material.

Here, we present a nuclear isolation protocol specifically optimized for long-term frozen brain tumor tissues and a possible workflow for identifying cell populations from snRNA-seq data. To study the usability of the nuclear preparations obtained with our protocol, we profiled the extracted nuclei using different snRNA-seq platforms: Chromium 10X Genomics^17^, Drop-seq (Macosko)^18^ and Fluidigm C1 System^19^. The resulting data were further analyzed computationally, showing the applicability of the derived data for identification of cellular subpopulations; assignment to cell types; copy number analysis and evaluation of lineage hierarchies.

## RESULTS

### Systematic optimization of nuclear isolation protocol for frozen tissues yields intact nuclei

We first tested the density gradient centrifugation method originally developed by Spalding et al^13^ and modified by Ernst et al^14^, and observed substantial cell debris and a low nuclear yield in the final preparation when using frozen human brain tumor tissues. In addition, we speculated that the extended processing time increased the risk of further RNA degradation during sample handling. In an attempt to improve nucleus yield and quality, we modified the respective buffers, optimized the extent of homogenization through douncing, added two filtering steps after the cell lysis and adapted the ultracentrifugation step. Despite these measures, we found that the sample purity was still insufficient (Fig. S1a). We therefore replaced the density gradient centrifugation with washing steps using lysis buffer without detergent, to avoid nuclear wall permeabilization. Although this clearly improved purity, the yield remained low. To overcome this, we tested several consumable plastics, coating buffers and buffer volumes and finally managed to increase the yield of nuclei (Fig. S1b; see Methods). The buffer volumes were adjusted based on the amount of starting material (here we present volumes suitable for about 20-50 mg of frozen glioma tissue).

Our final protocol is fast (less than 30 minutes in total), low-cost, and simple to carry out. It consists of four steps: cutting the tissue in ice-cold lysis buffer with a scalpel, douncing the sample to open cell walls, filtering the cell debris and washing off the rest of the cell debris and free RNA (Fig. 1a). The sample can then be re-suspended in storage buffer and either applied directly for use with snRNA-seq platforms or frozen for a short period (maximally a few days) at −80°C. Washing of the nuclei three times was found to be optimal for obtaining a debris-free supernatant (Fig. 1b), with further washing steps leading to damage of the nuclear walls. However, some nuclei are lost in each wash (Fig. 1c) and therefore two washes may be preferred if the amount of starting material is low. The protocol yields intact nuclei with undamaged nuclear walls in a debris-free supernatant (Fig. 1d).

**Figure 1.**
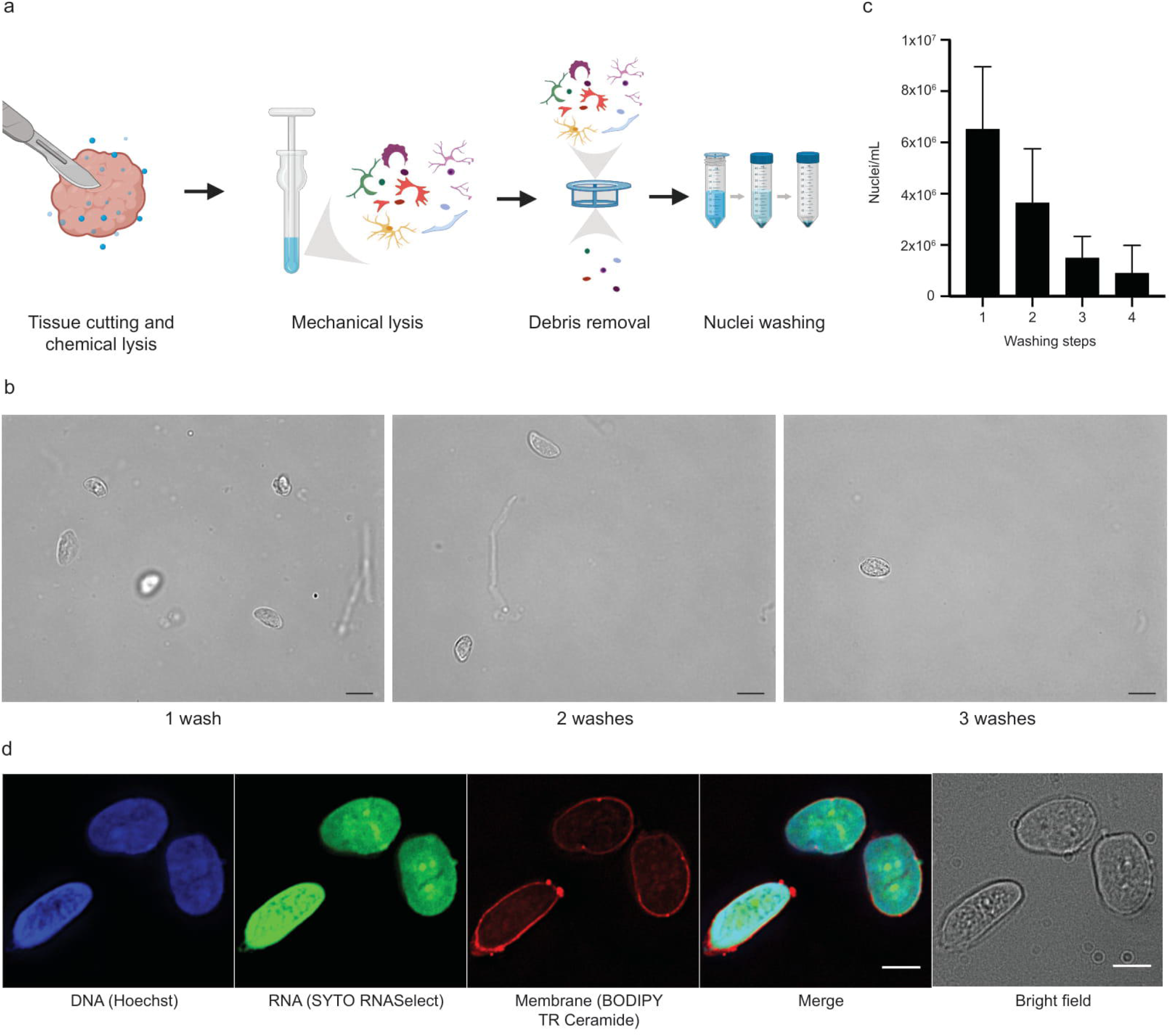
Isolation of intact nuclei from long-term frozen pediatric glioma tissue. (a) Schematic figure showing sample preparation steps. (b) Representative images showing the effect of an increasing number of washing steps on nuclei yield/integrity and amount of debris (scale bar 10 μm). (c) Nuclei yield decreases with increasing number of washes. (d) Staining for nuclear membrane, DNA and RNA reveals intact nuclei, with no leakage of nucleic acids (scale bars 5 μm). Nuclei originate from a pediatric pilocytic astrocytoma tissue frozen for seven years prior to nuclear extraction.

As a comparison, we also tested three commercial nuclear isolation methods: Nuclei EZ Prep (Sigma-Aldrich, NUC101-1KT), Isolation of Nuclei for Single-Cell RNA Sequencing (10X Genomics)^15^ and OptiPrep™^16^. The EZ prep resulted in a high yield, but also in a very high amount of debris in the supernatant (Fig. S1c), while the 10X Genomics protocol gave a clear supernatant but very low yield of nuclei (Fig. S1d). When replacing the washing step of our protocol with OptiPrep™ density gradient, we obtained similar results as when using our original sucrose cushion density gradient, but with a slightly lower yield (Fig. S1e). Our optimized protocol therefore provides a good balance between purity and yield, in addition to the benefits of its simplicity and cost-effectiveness.

### Comparison of snRNA-seq platforms

A pediatric glioblastoma patient-derived xenograft (PDX) sample (Table S1) was analyzed using Chromium (10X Genomics)^17^, Drop-seq (according to Macosko et al.)^18^ and C1 (Fluidigm)^19^ systems (Fig. S2a). As expected, the highest number of nuclei was detected with 10X Genomics, with Drop-seq detecting only slightly less. The lower throughput design of the plate based Fluidigm C1 platform generally produces a reduced number of analyzed nuclei (Table S2). Separation of mouse and human nuclei was clear with both droplet-based as well as plate-based methods, with few mixed signals (Fig. 2a, Fig. S2b-c). Although the Fluidigm C1 analysis resulted in a much higher detection of genes overall (~6000) compared with the droplet-based methods (10X Genomics ~2000 and Drop-seq ~1000) (Fig. 2b, Fig. S2d, Table S2), the proportion of human nuclei detected in comparison with mouse nuclei was greatly reduced (Fig. S2e). This difference may derive from physical differences of healthy mouse nuclei in comparison to tumorous human nuclei, and/or differences in nucleus size affecting capture efficiency, resulting in a differential flow in the C1 chip (although the results are based on small numbers overall). The poorer human to mouse nuclei ratio is also reflected in the presence of total human genes (Fig. S2f). Despite this difference, however, the most highly expressed genes were still similar between 10X and C1 (Fig. 2c, Fig. S2g-h). When comparing our snRNA-seq data of the PDX material merged as a pseudo-bulk with whole-tissue Affymetrix gene expression array data from multiple different patient tumors, the highest similarity was with the matched sample from which the PDX was derived (Fig. 2d). The other closely similar tumors belonged to the same tumor entity (pediatric high-grade glioma), with unrelated pediatric brain tumors showing a lower overall similarity. The same result was seen with both the 10X Genomics and the Fluidigm C1 datasets (not shown).

**Figure 2.**
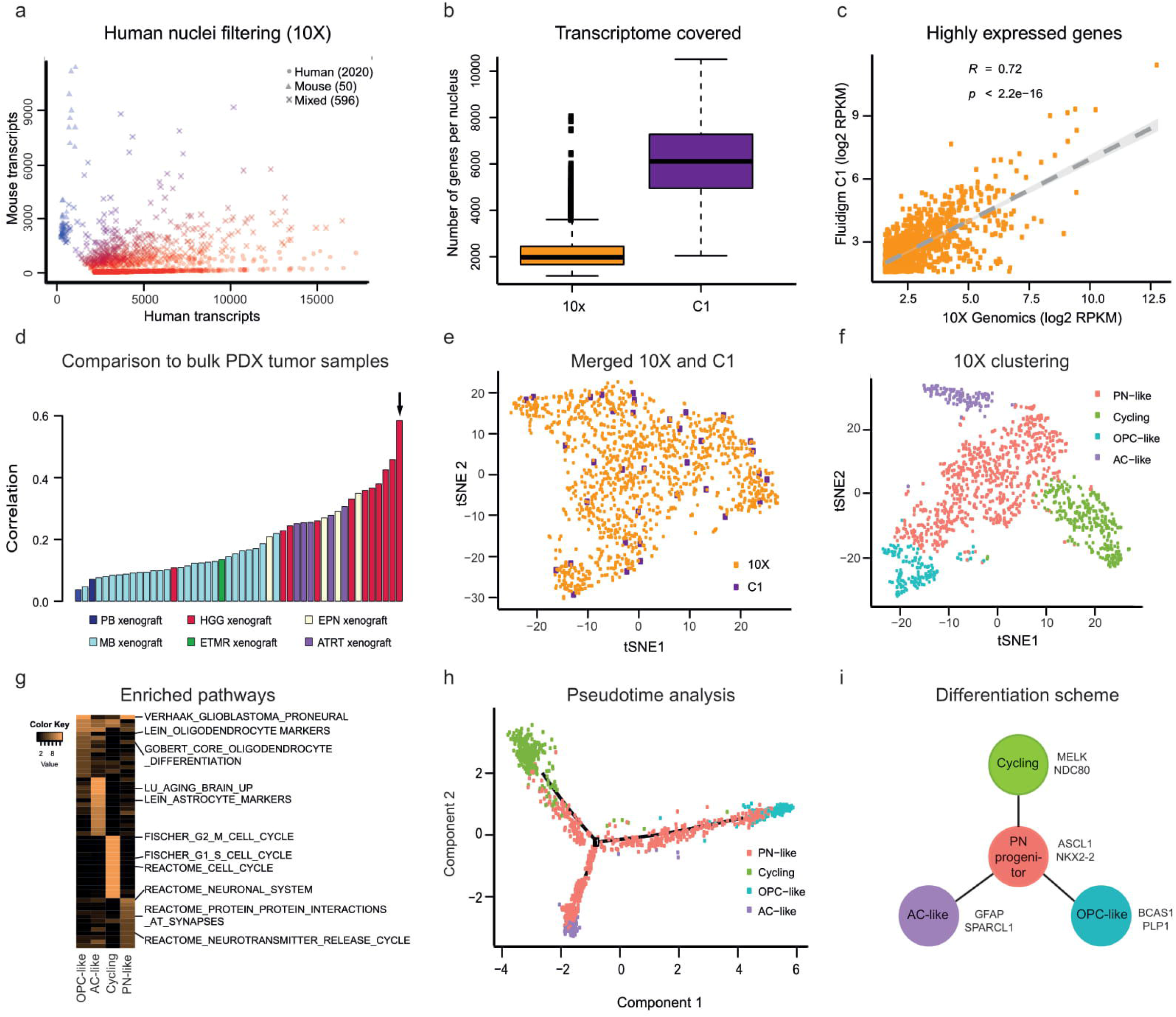
Chromium 10X Genomics is optimal for studying tumor heterogeneity using patient-derived xenografts. (a) Variance between the numbers of human and mouse transcripts per nuclei from 10X snRNA-seq data of a glioma PDX sample. (b) Number of genes per cell and (c) gene expression levels between 10X and C1 platforms. (d) Comparison of 10X PDX snRNA-seq data combined into pseudo-bulk to bulk microarray data from a group of PDX samples based on correlation, the same glioma PDX sample is marked with an arrow. (e) *t*-distributed stochastic neighbour embedding (t-SNE) representation of combined 10X and C1 scRNA-seq datasets. (f) t-SNE representation of 10X PDX dataset. (g) Heatmap of pathways enriched among 10X PDX cell types. Colors represent confidence level −log10 (p-val). (h) Pseudotime trajectory analysis of 10X PDX cells. (i) Schematic representation of the identified tumor cell populations.

Notably, more mitochondrial genes were found in the Drop-seq data compared with the two commercial platforms (Fig. S2i). The Fluidigm C1 platform has an additional washing step included, which might result in an improved filtering of the mitochondria. A clear explanation for the difference between the two droplet-based methods was not apparent, but one hypothesis could be that the lysis buffer in the Drop-seq may lyse the mitochondrial wall more efficiently than the lysis buffer used in 10X Genomics. The processing time of the Drop-seq is also higher than that for the 10X Genomics system, which results in a longer incubation time of the sample in the lysis buffer and thus possibly enhanced lysis of mitochondrial walls.

Although the inherent design differences of the two platforms means that more nuclei can be analyzed with 10X Genomics compared with Fluidigm C1, all of the identified cellular subpopulations were found in the data of both systems (Fig. 2e, Fig. S2j). Due to the clearer differences in cluster designation with increased nucleus number, however, all subsequent analyses were performed with data from the 10X platform.

The original clustering of the PDX sample results in seven suggested clusters (Fig. S2k). When assigning a putative identity to these clusters based on known marker gene expression (Fig. S2l, Table S3a) ^20^, we found evidence for four different cell populations: cycling cells, proneural-like (PN-like) cells, astrocyte-like (AC-like) cells, and oligodendrocyte precursor-like (OPC-like) cells (Fig. 2f). This assignment of the cell populations is further supported by analysis of enriched pathways (Fig. 2g, Table S4a). Pseudotime analysis of the nuclei data indicated three cellular states (Fig. S2m). Mapping of cellular identities onto the pseudotime trajectory based on expression of known marker genes (Fig. S2n), however, suggested a model in which a cycling subpopulation gives rise to an intermediate PN-like progenitor (consisting of mesenchymal-like and neural progenitor-like cells) that can subsequently differentiate into AC-like or OPC-like daughter cells (Fig. 2h-i). The identified cell states comprised distinct clusters on the original tSNE plot, as visualized by the expression levels of respective marker genes (Fig. S2o).

### Single-nucleus RNA-seq allows detailed molecular analysis of frozen primary human brain tumor samples

Nuclei from two long-term frozen primary pediatric glioma tissues (Table S1) were analyzed using the 10X snRNA-seq system, providing high-quality data with good confidence of transcriptome mapping (Table S2). The samples were sequenced on three different machines (Illumina NextSeq500, HiSeq4000 and NovaSeq6000), with the estimated number of nuclei detected not substantially differing between the platforms (Table S2). As expected based on their output, the sequencing saturation with NextSeq500 was lower than for the HiSeq and NovaSeq (estimated saturation of detected genes/nuclei was at approximately 100,000 reads per nucleus). This could be compensated for by reduced pooling of samples, and the proportion of reads mapped to the genome/transcriptome showed that the general performance was at least as good with the NextSeq500 as with the other two devices.

The first tumor examined was a pediatric high-grade glioma (HGG), one of the most lethal forms of pediatric brain tumor with a typical survival of less than 2 years after diagnosis^21^. Our extraction protocol generated a high-yield of clean nuclei for sequencing from the frozen tissue specimen (Fig. 3a). Analysis of the single nuclei data suggested two predominant clusters. Based on previously reported marker genes for glioblastoma (GBM) tumor cell populations^20^, the major cluster in this sample (ICGC_GBM61, as reported in ^22^) seems to consist of neural progenitor cell-like (NPC-like) tumor cells (Fig. 3b). An additional second cluster was also observed. Based on CHETAH^36^ cell type classification using healthy brain cell types as reference, this cluster demonstrated high similarity to microglia (Fig. 3c). Further evaluation of marker genes for the tumor and microglial nuclei confirmed the discrete nature of the clusters (Fig. 3d-f).

**Figure 3.**
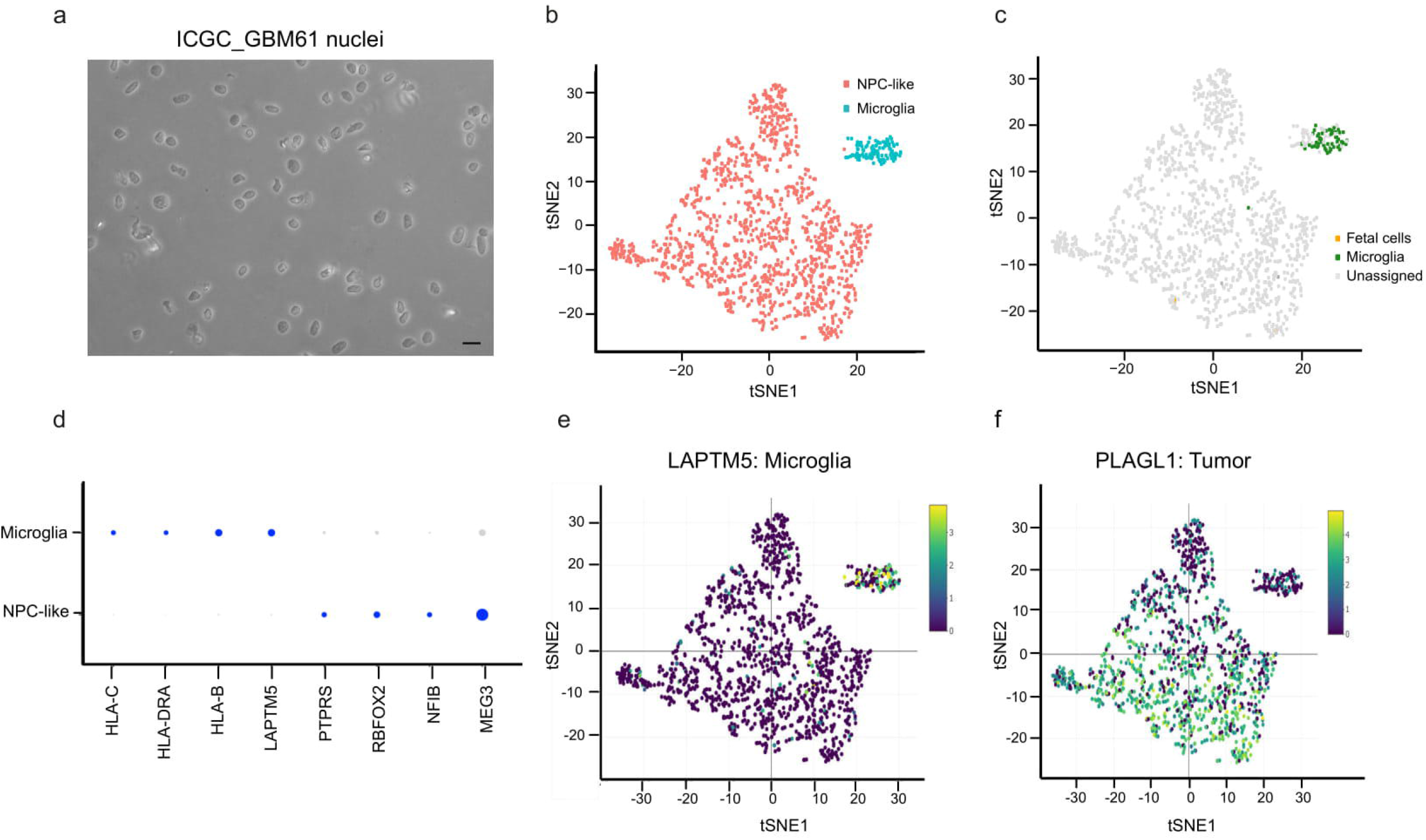
snRNA-seq of ICGC_GBM61 reveals distinct tumor and healthy cell populations. (a) Brightfield image showing intact isolated nuclei loaded to the 10X system (scale bar 10 μm). (b) t-SNE representation of ICGC_GBM61 snRNA-seq data. Neural progenitor cell-like (NPC-like) tumor cells and microglia form separate clusters. (c) CHEETAH confirms the identity of a distinct microglia cluster. (d) The most highly expressed marker genes show differential expression across assigned cell types. Expression of (e) LAPTM5 representing microglia and (f) PLAGL1 in tumor cells.

Tools identifying tumor cell populations based on copy number variations (CNVs) have been developed for whole-transcriptome scRNA-seq methods such as the Fluidigm C1 or Smart-seq2 approaches^23,24^. The 10X Genomics system, however, amplifies only the 3’ ends of transcripts during preparation of the cDNA library. The total number of transcripts detected is also typically lower, making expression-based estimates of genomic copy number challenging. There are, however, methods being developed that also appear to be applicable to 10X data. One such tool for assessing copy number changes in tumor cell clusters from 10X snRNA-seq, InferCNV^8^, gave promising results despite not being optimised for 3’ read data (Fig. 4a). The healthy microglia cluster of ICGC_GBM61 was used as an internal reference for comparison with the tumor clusters of the same sample, with clear indications for regions of copy number change observed. While the overlap with the bulk CNV plot of the same tumor was limited overall, combining the single nucleus CNV results into pseudo-bulk demonstrated a reasonable correspondence for certain regions (Fig. 4b). For example, the calling of loss of chromosome 16, gain on chromosome 6q and MYCN amplification could potentially be used as an additional confirmation to annotate any ambiguous clusters as representing either tumor or stromal cells.

**Figure 4.**
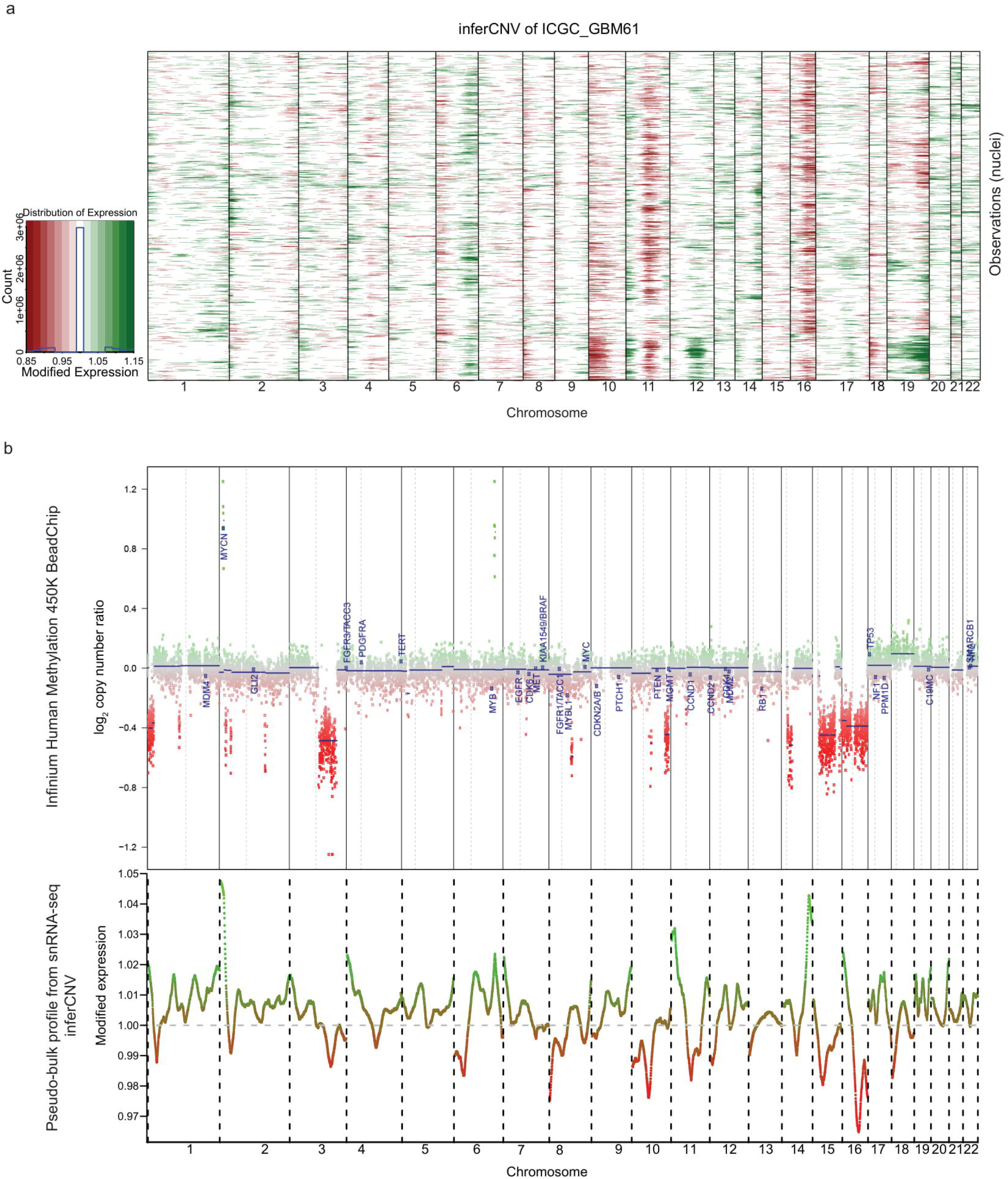
Copy number variation analysis supports tumor cell detection from 10X snRNA-seq data. (a) Copy number variations (CNVs) of single nuclei from 10X snRNA-seq data of ICGC_GBM61 analyzed by inferCNV. (b) A bulk CNV profile of the same tumor derived from Infinium HumanMethylation450 array analysis compared to pseudo-bulk extraction of CNV profiles from 10X data of ICGC_GBM61 using mean values across the cells.

We also profiled a pilocytic astrocytoma sample (PA, a WHO grade I tumor representing the most common childhood brain tumor) using snRNA-seq, again with multiple sequencing devices. The nuclei preparation was again of good quality, despite a tissue storage time of 6 years (Fig. 5a). Overall cluster detection was similar with each of the sequencers (Fig. S3a-c), with some minor differences in cluster assignment of individual nuclei between platforms (Fig. S3d). The clearest picture was obtained when combining all of the data, confirming that a sufficient overall sequencing depth is more important for the identification of distinct cell populations than the specific device used to generate the data (Fig. S3e).

**Figure 5.**
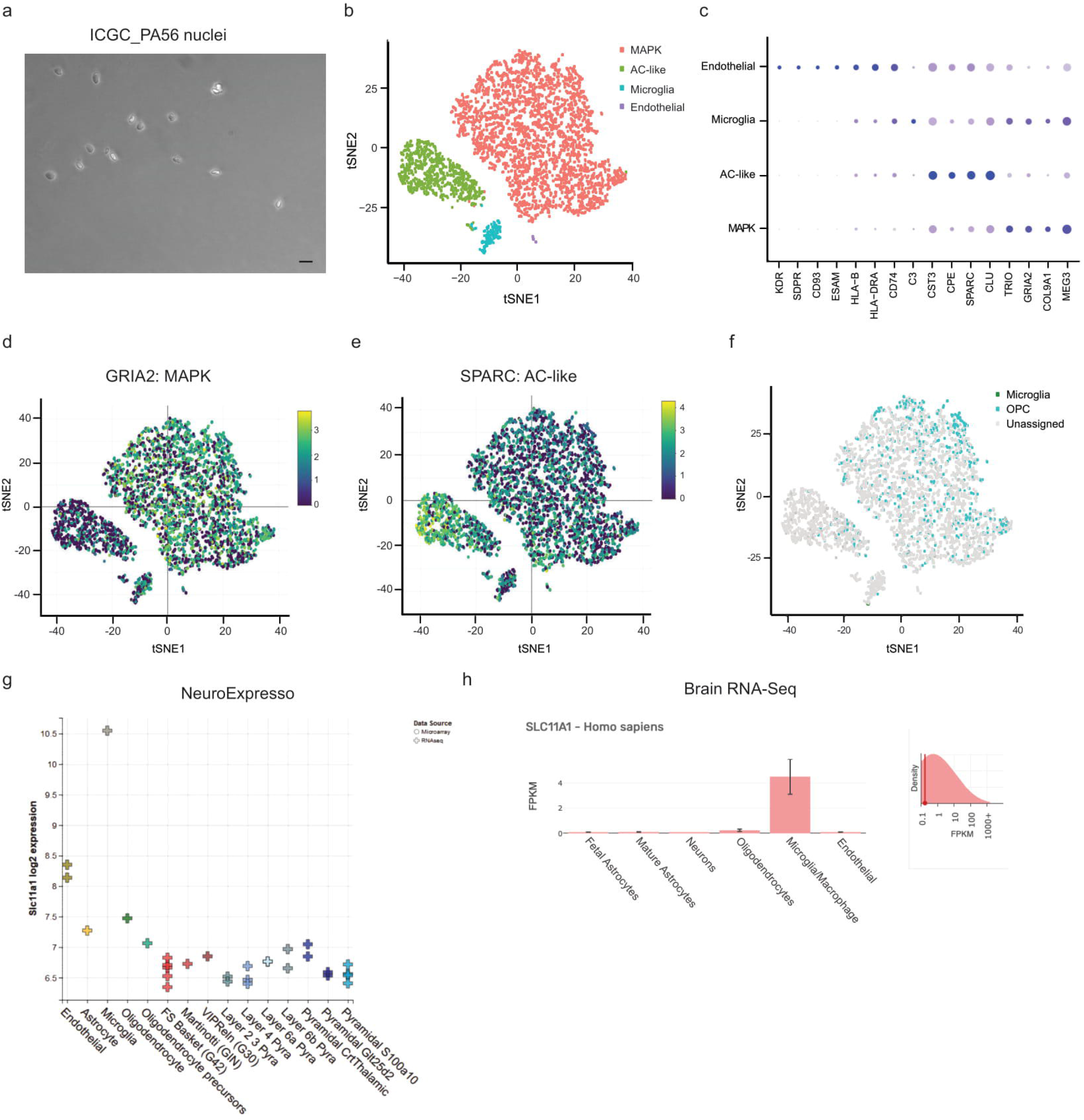
Combining several tools helps to assign an identity to unknown cell populations. (a) Representative image of the isolated nuclei from ICGC_PA56 (scale bar 10 μm). (b) t-SNE representation of the PA dataset. (c) Representation of highly expressed marker genes within the cell clusters representing AC-like tumor cells, MAPK program expressing tumor cells, microglia and endothelial cells. Expression of genes representing (d) MAPK pathway activation and (e) AC-like cells. (f) CHEETAH analysis using a healthy brain reference reveals OPC-like cells in the tumor cell cluster of ICGC_PA56. (g) NeuroExpresso and (h) Brain RNA-Seq databases confirm the identity of the putative microglia cluster.

When looking in detail at this tumor (ICGC_PA56, harboring a KIAA1549:BRAF fusion – the most common genetic alteration in this entity^25^), a number of distinct cell types could be observed (Fig. 5b). Based on previously published marker genes for PAs^28^, the largest tumor cell cluster was found to express genes associated with an active MAPK pathway (Fig. 5b-d), confirming the key role of this pathway in PA^26^. The other major cell population in ICGC_PA56 was identified to be AC-like tumor cells based on marker gene expression (Fig. 5e). These cells did not, however, resemble normal, healthy astrocytes based on CHETAH analysis or when looking at highly expressed genes (Table S3) using NeuroExpresso^40^ and Brain RNA-seq^39^ database (Table S5c), thus indicating that they most likely represent a second (more differentiated) tumor cell subpopulation. When comparing with normal brain cells, many of the single tumor nuclei showed similarities with oligodendrocyte precursor cells (OPCs; Fig. 5f), which have previously been suggested to be the cells-of-origin of PA tumors^27^. This similarity was particularly notable in the MAPK subset, further supporting a key role for MAPK-activated, OPC-derived tumor cells as a main component of PAs.

Two putative healthy cell populations were also identified within the PA sample (Fig. 5b). In contrast to the tumor cells, which often gave no clear match, these populations give a relatively uniform result when comparing with NeuroExpresso and the Brain RNA-seq database as matching to microglia (Table S5b-c). These tools thus proved valuable for confirming the identity of clusters that may be otherwise difficult to assign (Fig. 5g-h). When combining the snRNA-seq data of the low- and high-grade glioma samples, the non-tumor cells in common between the two were found to cluster more closely to each other than to other cells from the respective tumors, while the two tumor clusters were clearly distinct (Fig. S4a). This combined analysis suggested a small fraction of endothelial cells in the HGG sample, which was not identified in the single tumor analysis, and confirmed the presence of microglia in both tumors. Despite their overall similarities, however, the immune cells also showed some differences (Fig. S4b), possibly hinting at different types or activation states of the microglia present. Further examination of these differences in future may provide information on, for example, tumor-promoting versus tumor-suppressing immune programs.

## Discussion

Here we have developed a method for extracting intact single nuclei from long-term frozen tissues. The method is simple, fast, low-cost, and is suitable for most typically equipped biology laboratories. The short processing time potentially reduces the risk of RNA degradation compared with some other protocols, and the method results in nuclei preparation with a good balance between purity and yield.

We found that nuclei extracted with this method can be applied to different scRNA-seq platforms. Both, a full transcriptome amplification method (Fluidigm C1) and a 3’ amplification method (10X Genomics) proved suitable for the analysis of nuclei isolated from frozen glioma tissues, with minor differences in results deriving from general design characteristics of the platforms. The 3’ transcriptome amplification method, for example, detected fewer genes per nuclei (although this could also be partly linked to lower read-count per cell, and will likely improve with upcoming chemistry developments) but more nuclei in total, which lead to more distinct clustering and a more accurate distribution of human and mouse cell proportions. Due to the inherent design of the 3’ amplification method, genomic mutations could not be detected, but InferCNV analysis did allow a broad detection of copy number variations.

Our study supports previous findings^11,28,29^ that nuclei from frozen tissues are a robust input material for single-cell omics analyses, thereby extending the range of suitable amenable to these high-resolution profiling techniques. This also removes need to sort for viable whole cells, which possibly reduces sources of technical variation in the analyzed tissues. A bias for certain cell populations based on differential sensitivity to the mechanical forces or lysis buffers applied in this protocol, however, cannot be excluded at present. Thus, it must be kept in mind that the detected nuclei populations might differ quantitatively (and possibly also qualitatively) from the original cell composition.

Applying this method to pediatric brain tumor tissues also showed its potential for revealing biological insights. We were able to detect distinct tumor cell subpopulations in both a low- and high-grade glioma sample, and also identified an infiltrating microglia component in both tumors. The application to PDX samples may also be of interest in future, for example for unambiguously examining contributions of tumor (human) vs stromal (mouse) cell types to the overall signaling milieu in the bulk tissue.

In summary, the possibility to extract and use nuclei from long-term frozen tissue material with a quick and simple protocol opens up enormous resources for the study of tumor heterogeneity and other questions of biological interest, expanding the utility of this rapidly developing method.

## METHODS

### Optimized nuclear extraction from frozen brain tumor tissues

All surfaces were cleaned with RNase Zap (Invitrogen AM9780) and PCR Clean wipes (Minerva Biolabs 15-2001) prior to sample processing, and equipment were cooled on ice and coated (0.1% Triton X-100 [Sigma-Aldrich 93443] in filtered PBS (Merck Millipore SLGS033SS and Gibco 14190-094). Only pipette tips of 1 ml (Biozym Low binding SafeSeal tips) and smaller were used to avoid losing the nuclei on the pipette tip walls. A fresh frozen human brain tumor tissue piece (20-50 mg) was placed on a Petri dish (Greiner Bio-One 628160) on a cooled metal block. It was mechanically dissociated using a scalpel in 1 ml of lysis buffer (0.32 M sucrose [Sigma-Aldrich 84097], 5 mM calcium dichloride [Sigma-Aldrich 21115], 3 mM magnesium acetate [Sigma-Aldrich 63052], 2.0 mM EDTA [Invitrogen 15575-038], 0.5 mM EGTA [Alfa Aesar J61721], 10 mM Tris-HCl, pH 8.0 [Invitrogen AM98556], 1 mM DTT [Sigma-Aldrich 10197777001] and 0.1% Triton X-100 [Sigma-Aldrich 93443]). An additional 4ml of lysis buffer were added, and the tissue was dissociated further by pipetting and then transferred into a glass douncer (Sigma-Aldrich D9063). The cells were mechanically lysed by douncing 10 strokes with pestle A and then 10 strokes with pestle B. The lysate was directly filtered using a 100 μm filter (Greiner Bio-One 542000) followed by a 40 μm filter (Greiner Bio-One 542040) to remove the bigger cell membrane debris and then spun down to remove the lysis buffer (500 g, 5 min, 4°C). The nuclei were re-suspended in 5 ml washing buffer (0.32 M sucrose [Sigma-Aldrich 84097], 5 mM calcium dichloride [Sigma-Aldrich 21115], 3 mM magnesium acetate [Sigma-Aldrich 63052], 2.0 mM EDTA [Invitrogen 15575-038], 0.5 mM EGTA [Alfa Aesar J61721] and 10 mM Tris-HCl, pH 8.0 [Invitrogen AM98556]) in a 50 ml tube (Corning 352070) (when having a frozen cell pellet as starting material, 3 ml of washing buffer and 15 ml tubes [Greiner Bio-One 188-271] were used). The debris and leaking RNA were removed by centrifugation (500 g, 5 min, 4°C). The washing steps should be done optimally two to three times, or a maximum of four times if extensive debris is observed (more washing rounds result in broken nuclear membranes). After the washing steps the nuclear pellet was re-suspended in 1 ml of nuclei storage buffer (0.43 M sucrose [Sigma-Aldrich 84097], 70 mM potassium chloride [ThermoFischer Scientific, AM9640G], 2 mM magnesium dichloride [ThermoFischer Scientific AM95306], 10 mM Tris-HCl, pH 7.2 [Sigma-Aldrich T2069] and 5 mM EGTA [Alfa Aesar J61721]). The nuclei can be processed directly in snRNA-seq platforms in the storage buffer or frozen shortly (up to ~1 week) at −80°C.

### Comparison to other nuclear extraction methods

The final method was developed based on the density gradient centrifugation protocol originally developed by Spalding et al 2005^13^ and modified by Ernst et al 2014^14^. The optimized protocol was compared to three commercial methods in addition to the original density gradient centrifugation method mentioned above using 40 mg of a frozen pilocytic astrocytoma tissue in each (Fig. S1). The commercial methods were Nuclei EZ Prep (Sigma-Aldrich, NUC101-1KT), Isolation of Nuclei for Single-Cell RNA Sequencing (10X Genomics)^15^ and OptiPrep™^16^ and were carried out following the manufacturer’s instructions. OptiPrep™ was used as an alternative density gradient centrifugation method and applied after cell lysis and filtering. The nuclear preparations were observed for quality using light microscopy.

### Nuclei staining experiments and microscopy

To test the optimal number of washing steps, 37 mg each of three different frozen glioma samples were processed using the optimized nuclear isolation protocol. Each sample was washed four times after nuclear isolation. After each wash, the nuclear pellet was re-suspended in 1 ml of storage buffer and a 50 μl sample taken. A further 4ml of washing buffer was added before continuing to the next washing step. The nuclei were counted using an automated cell counter (Fig. 1c) and imaged using a light microscope (Zeiss Cell Observer) (Fig. 1b).

To study the condition of the isolated nuclei and possible RNA leakage, the nuclei were stained using DNA, RNA and membrane stains. Nuclei were isolated from frozen pediatric pilocytic astrocytoma tumor tissue using the optimized protocol with three washing steps. The nuclei were stored at −80°C overnight. On the next day, the nuclei were stained with RNA stain (SYTO™ RNASelect™ Green Fluorescent cell Stain [ThermoFischer Scientific, S32703]), DNA stain (Hoechst 33342 Solution [ThermoFischer Scientific, 62249]) and membrane stain (BODIPY™ TR Ceramide [ThermoFischer Scientific, D7540]) and imaged using a fluorescent microscope (Olympus IX71) (Fig. 1d).

### snRNA-seq experiments

Libraries from single nuclei were generated using either Chromium 10X Genomics^17^, Fluidigm C1^19^ or Drop-seq^18^ platforms according to the manufacturers’ instructions as described in Macosko et al^18^ and Bageritz et al^30^ (for Drop-seq). Nuclei were loaded in a concentration of 300 nuclei/μl into 10X and C1 and 275 nuclei/μl into Drop-seq. For Drop-seq, nuclei concentration was adjusted to the smaller droplets to not show more than 5 % doublets. The PDX sample was loaded freshly after nuclear isolation into each snRNA-seq system, and the primary human tumor nuclei were frozen overnight at −80°C prior to 10X experiments. Single Cell 3’ Reagent Kit v2 was used for 10X while SMARTer Ultra Low RNA Kit for the Fluidigm C1 System (small-cell IFCs). After the cDNA library construction, all the 10X samples were sequenced on a HiSeq4000 sequencer (Illumina), and some of the samples (Table S2) using NextSeq 500 (Illumina) or NovaSeq6000 (Illumina) (paired end 26+74 bp). The C1 libraries were sequenced on a HiSeq2000 (50 bp single-end dual index reads, 2×8 bp) and Drop-seq libraries on a HiSeq2500 (paired end 20+180 bp, 8 bp index).

### Bulk sequencing experiments

RNA from frozen tumor bulk tissues was used for Affymetrix gene expression array (Affymetrix Human Genome U133 Plus 2.0) and DNA in Infinium Methylation EPIC kit (Illumina).

### Ethical justification

The PDX mouse used in this study was handled in accordance with legal and ethical regulations and approved by the regional council (Regierungspräsidium Karlsruhe, Germany; G-64/14). Informed consent was obtained according to ICGC guidelines for all of the human tumor tissues used in this study.

### snRNA-seq data analysis

The initial processing of sequencing reads was performed independently for each platform. For 10X data the original CellRanger v2.1.0 pipeline was applied with an intron-including genome reference. Fluidigm C1 read alignment was performed with STAR v2.5.2b^31^ to the genome reference combined with ERCC and gene expression counts computed using featureCounts v1.4.6^32^. Drop-seq reads were processed using Drop-seq tools v1.12 ^18^. For the alignment of PDX samples, combined hg19/mm10 reference was used, while tumor sample processing was performed with hg19 only. For PDX samples the differentiation between human and mouse cells was performed based on the comparison of transcript counts per species. The filtering cut limit for each platform was selected based on computation of the mean proportion of human genes per cell.

Quality control was performed for each sample using Scater package v1.8.0^33^. The cell clustering for 10X data was generated using the Seurat v2.3.2 package^34^. The integration of 10X and C1 data for the PDX sample was performed using Canonical Correlation Analysis (CCA) from Seurat. Trajectory reconstruction was achieved from the application of Monocle v2.8.0^35^ on the adjusted result from Seurat. Correspondence of the cell types to specific pathways was computed using hyper geometric test applied on functional gene lists from the MSigDB collection^36^. Combined visualization of cluster similarity between platforms was performed using GoogleVis R package with assignment based on a positive correlation limit of 0.3.

CHEETAH v1.0.4 R package^37^ was applied for the comparison of our single nucleus data to other reference datasets. A reference for healthy brain cell types was acquired from the study of Darmanis et al^38^ while the reference data set for the tumor cells (H3^K27M^ mutated pediatric high grade glioma) was from the study of Filbin et al^7^. Default confidence threshold (0.1) was used in the analysis. Copy number profiling was performed from the usage of inferCNV v1.3.2 of the Trinity CTAT Project as previously described^5^. In addition, two online databases, Brain RNA-Seq^39^ and NeuroExpresso^40^, were used for studying the differentially expressed genes in each cluster. Gene expression levels per nucleus were visualized using a custom analysis script kindly made available by Dr. Murat Iskar (unpublished).

### Affymetrix data integration

The Affymetrix PDX data was collected from Brabetz et al. ^41^. Mean values between probes per gene were computed to generate a full normalized expression matrix. PDX single nucleus profiles (10X and C1) were combined into bulk datasets and normalized via RPKM for the comparison with Affymetrix data. Correlation was computed based on the selection of the top 500 most highly variable genes in common between the snSeq sample and the Affymetrix matrix.

## Acknowledgements

We thank Katharina Bauer for help and guidance in snRNA-seq experiments, and Thorsten Kolb and Aurélie Ernst for their advide with the sucrose gradient-based nuclear isolation protocol. We further thank Henrik Kaessmann and Mari Sepp for their ideas regarding the development of the isolation protocol, Sebastian Brabetz and Norman Mack for providing the PDX sample and Murat Iskar for developing the graphical user interface tool for visualization of gene expression in single nuclei. Support by the German Cancer Research Center (DKFZ) Single-Cell Open Lab (scOpenLab), Genomics & Proteomics Core Facility as well as Light Microscopy Facility is gratefully acknowledged. The study was supported by funding of the Everest Centre for Low-Grade Paediatric Brain Tumour Research (The Brain Tumour Charity, UK; GN-000382).

## Author contribution

Conceived the idea and supervised the project: M.B., S.M.P., D.T.W.J., M.Z.; performed the experiments: K.J.E., J.B., J-P.M., A.W., K.K.M., S.L.; performed bioinformatics analyses: K.O., K.J.E; wrote the manuscript: K.J.E., K.O., D.T.W.J., M.Z., S.M.P. All authors approved the final manuscript.

---

**Supplementary Figure 1.**
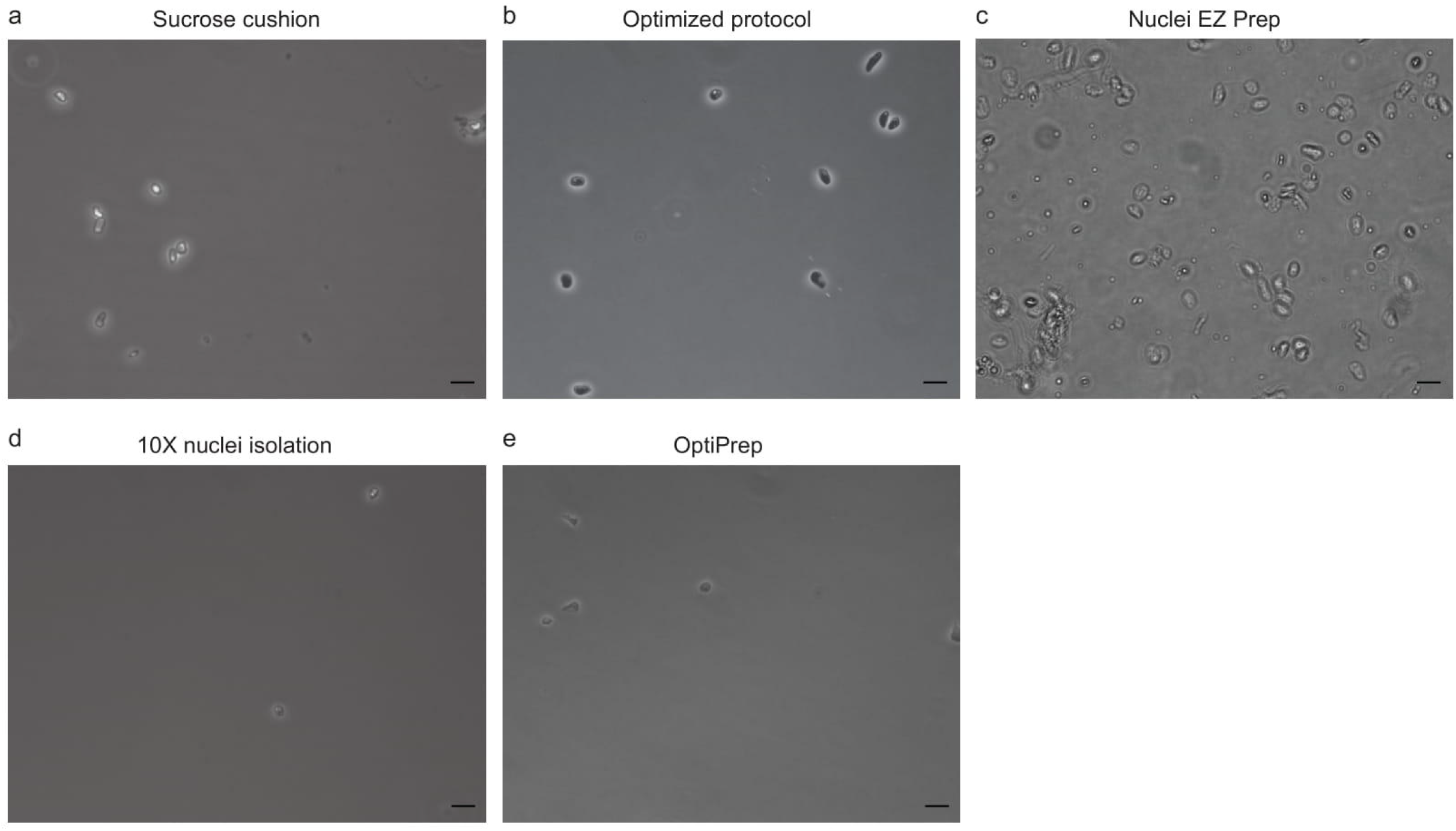
Comparison of the optimized nuclei isolation protocol to other protocols. Brightfield image of nuclei extracted from a pilocytic astrocytoma tissue using (a) sucrose cushion density gradient, (b) our optimized protocol, (c) Nuclei EZ Prep, (d) Isolation of Nuclei for Single-Cell RNA Sequencing (10X Genomics) and (e) OptiPrep™ (scale bar 10 μm).

**Supplementary Figure 2.**
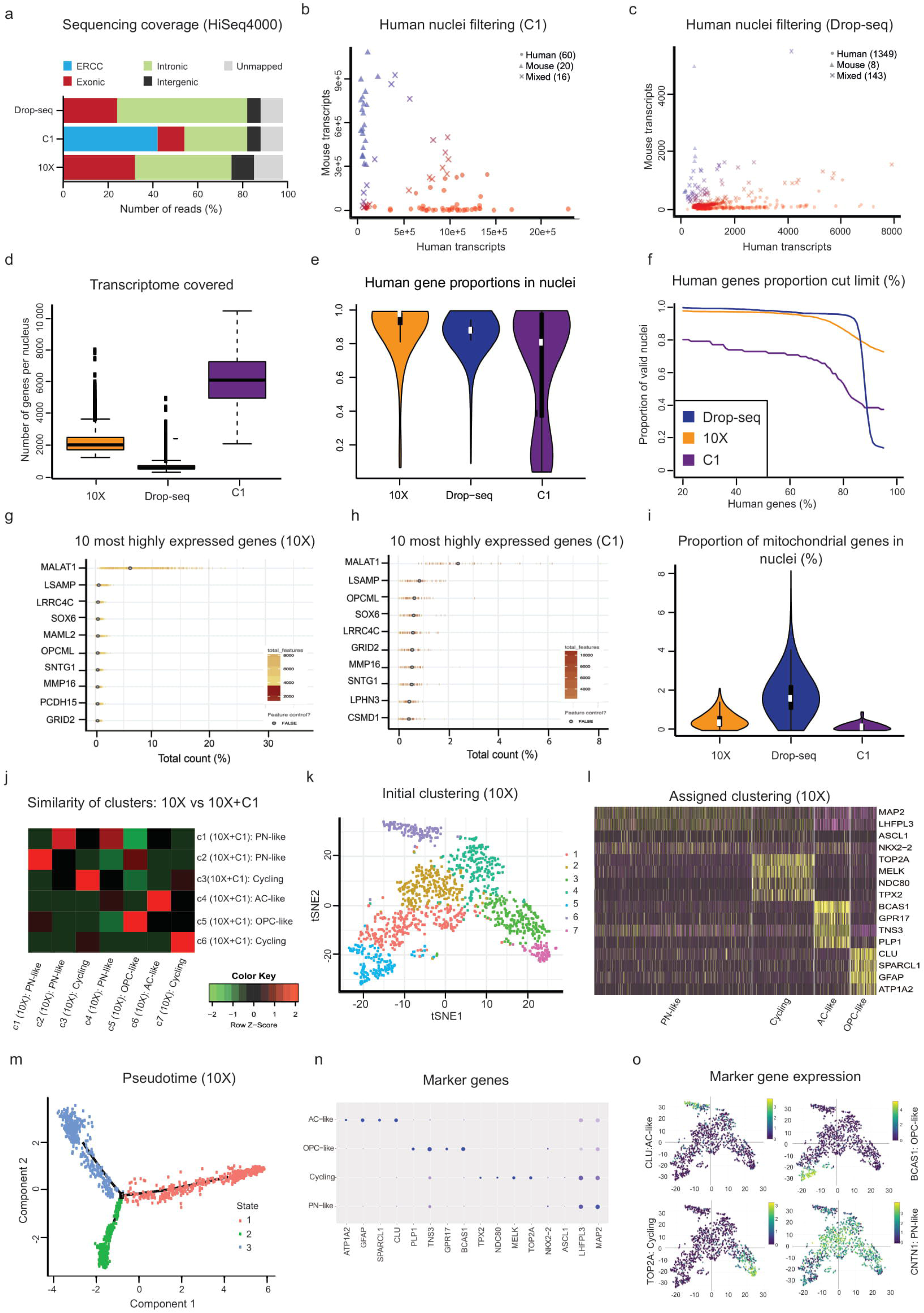
Comparison of 10X Genomics, Fluidigm C1 and Drop-seq using a patient-derived xenograft sample shows advantage for 10X Genomics due to high number of nuclei detected. (a) Proportions of covered reference types from reads alignment of three different snRNA-seq platforms are fairly similar, with the exception of ERCC spike-ins with Fluidigm C1 data. (b,c) The numbers of human and mouse transcripts per nuclei from of a glioma PDX sample for (b) Fluidigm C1 and (c) Drop-seq data. (d) Boxplot represents numbers of total genes per nuclei between 10X, Drop-seq and C1 platforms. (e) The violin plot represents proprotions of human materials per nuclei among platforms. Proprotions are computed as a ratio between numbers of human only and total transcripts in a nuclei. (f) Effect of the minimum proprotion of human genes cut limit to assign nuclei as valid (not mixed or from mice). Most highly expressed genes were similar in (g) 10X and (h) C1. (i) Proportions of mitochondrial genes per nuclei between 10X, Drop-seq and C1. (j) Correlation-based comparison of detected clusters between 10X and 10X+C1 combined. (k) t-SNE representation of 10X PDX dateset, original clusters are marked in color. (l) Heatmap of four main assigned cell type specific differentially expressed genes. (m) Original pseudotime trajectory plot before assigning the populations based on (n) high expression of known marker genes (Fig. 2h). (o) Visualization of the assigned cell type marker gene expression in t-SNE representation of 10X PDX.

**Supplementary Figure 3.**
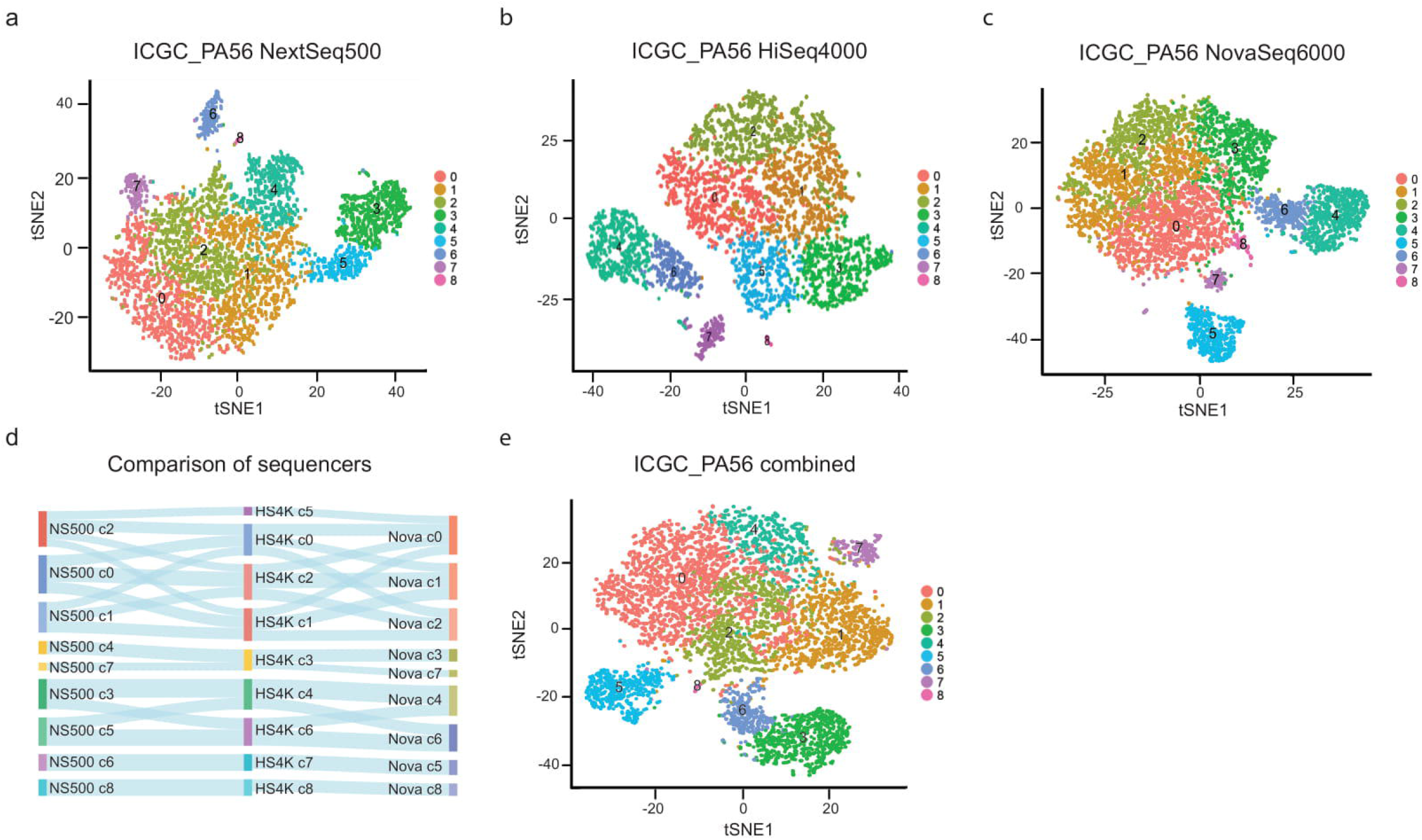
HiSeq4000, NextSeq500 and NovaSeq6000 provide corresponding cluster detection. t-SNE representation of ICGC_PA56 10X snRNA-seq data sequenced with (a) NextSeq500, (b) HiSeq4000 and (c) NovaSeq6000 resulting. (d) Assocations between the cell clusters of platforms based on positive correlation limit 0.3 (e) t-SNE representation of ICGC_PA56 snRNA-seq dataset combined from all platforms.

**Supplementary Figure 4.**
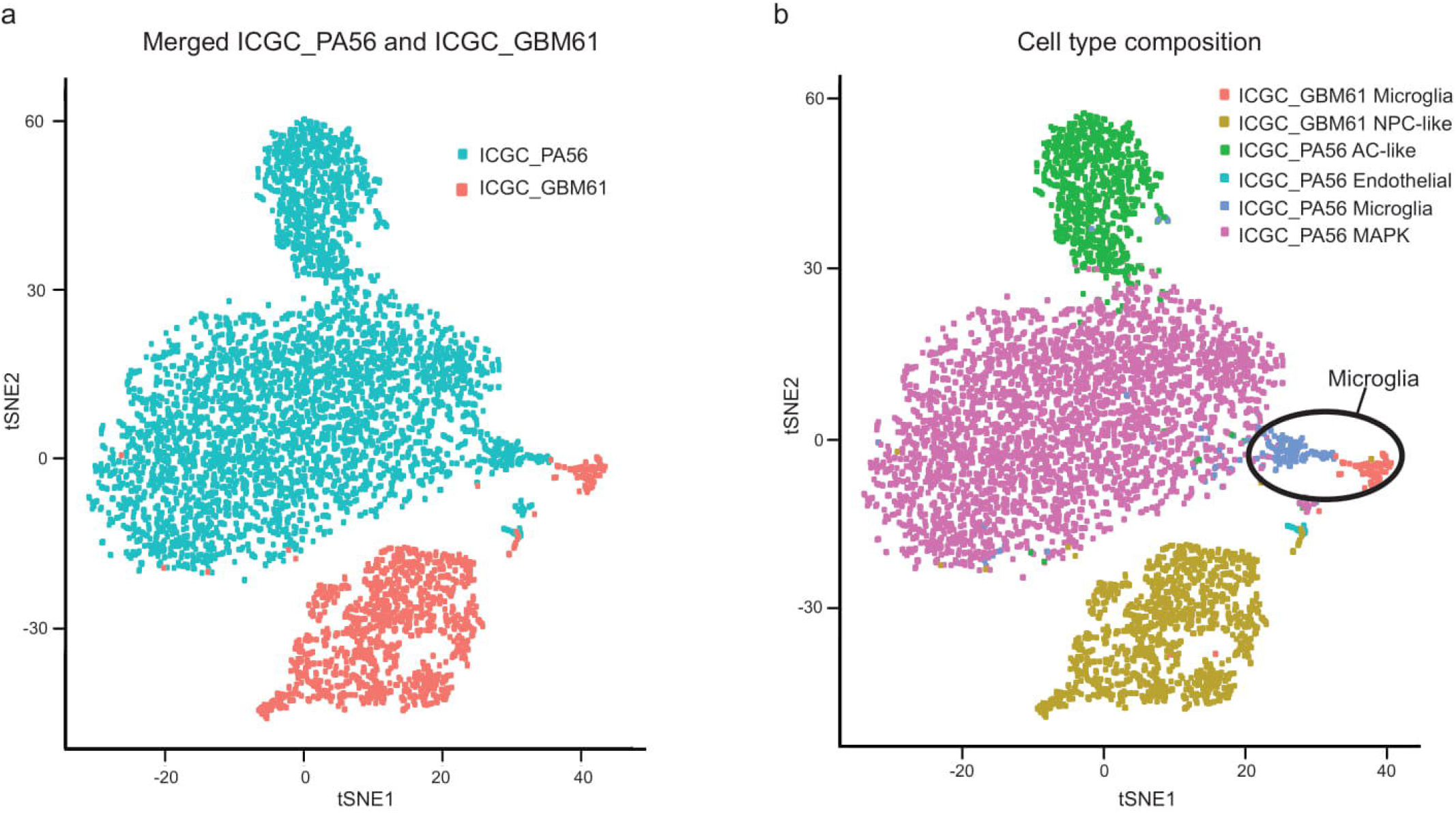
Tumor cell populations from different snRNA-seq tumors cluster more distinctly, while healthy populations are more proximal to each other. (a) ICGC_PA56 and ICGC_GBM61 10X snRNA-seq populations cluster separately as observed in a t-SNE plot. (b) Healthy normal microglia cells (marked in color) from both tumors cluster close to each other.

## Supplementary Tables

**Supplementary Table 1.** Clinical chacteristics and sample processing information of the pediatric glioma tissues studied by snRNA-seq.

**Supplementary Table 2.** Quality control information of the pediatric glioma tissues and the PDX model studied by snRNA-seq.

**Supplementary Table 3.** Upregulated genes per each cluster

**Supplementary Table 4.** Enrichment analysis of specific upregulated DEGs based on GSEA MSigDB 7.0 database using hypergeometric test.

**Supplementary Table 5.** 10 most gighly upregulated genes per each cluster assigned to cell types based on NeuroExpresso and Brain-RNA-seq databases.

